# Cassava planting material movement and grower behaviour in Zambia: implications for disease management

**DOI:** 10.1101/528851

**Authors:** Anna Maria Szyniszewska, Patrick Chiza Chikoti, Mathias Tembo, Rabson Mulenga, Christopher Aidan Gilligan, Frank van den Bosch, Christopher Finn McQuaid

## Abstract

Cassava is an important food crop for most small-holder growers across sub-Saharan Africa, where production is largely limited by the presence of two viral diseases: cassava mosaic disease (CMD) and cassava brown streak disease (CBSD), both propagated by a vector whitefly and via human-mediated movement of infected cassava stems. Despite its importance, there is limited knowledge of growers’ behaviour related to planting material movement, as well as growers’ perception and knowledge of cassava diseases, which have major implications for disease spread and control. This study was conducted to address the knowledge gaps by surveying small-holder growers in Zambia. A total of 96 subsistence cassava growers across five provinces were surveyed between 2015 and 2017. Most growers interviewed used planting materials from their own (94%) or nearby (<10 km) fields of family and friends, although some large transactions with markets, middlemen, and NGOs occurred over longer distances. Information related to cassava diseases and uninfected planting material, however, only reached 48% of growers. Growers with access to information were more concerned about the disease, compared to uninformed growers. These data provide a basis for future planning of cassava clean seed systems to control virus diseases, emphasising the critical role of grower knowledge, and consequently education, in success of these systems. In particular, we highlight the importance of extension workers in this education process, as well as farmer’s groups and the media.

## Introduction

Cassava (*Manihot esculenta* Crantz) is one of the most important root crops in Zambia, and is a staple consumed throughout the year in Western, North Western, Luapula, and Northern provinces. Despite the importance of the crop, Zambia suffers from very low average national yields of 5.8 tonnes per hectare (t/ha) (FAOSTAT 2018). This is considerably lower than the reported average yield of neighbouring countries: Malawi (22 t/ha), Angola (10.9 t/ha) and Democratic Republic of Congo (8.1 t/ha) (FAOSTAT 2018). The low yield is in part due to the high prevalence in most of the cassava-growing areas of cassava mosaic disease (CMD, caused by cassava mosaic geminiviruses family *Geminiviridae*, genus *Begomovirus*) (Chikoti et al. 2013). This disease is the most prevalent and devastating disease of cassava in sub-Saharan Africa, causing considerable losses in yield (Legg et al. 2006; Muimba-Kankolongo et al. 1997; Szyniszewska et al. 2017; Thresh et al. 1997). To make matters worse, in 2017 cassava brown streak disease (CBSD, caused by potyviruses, family *potyviridae*, genus *Ipomovirus*), was confirmed in both Northern and Luapula provinces of Zambia (Mulenga et al. 2018). Both diseases are transmitted by the whitefly vector *Bemisia tabaci* (order *Hemiptera*, family *Aleyrodidae*), and through human-mediated vegetative propagation of infected planting material (Maruthi et al. 2017). Both CMD and CBSD are of great concern across sub-Saharan Africa because of their detrimental impact on root yield and quality (Abaca et al. 2012; Alvarez et al. 2012; Mbanzibwa et al. 2011; Winter et al. 2010). Both diseases increase poverty by dramatic loss in yield, and continue to deteriorate the livelihoods of millions of Africans (Legg and Thresh 2003; Patil et al. 2015).

Strategies for disease mitigation include the removal of infected plants (rouging), the adoption of resistant cultivars, and the use of disease-free planting material (known as “clean seed”). Each strategy faces particular challenges; difficulty in identifying infected plants, a paucity of resistant varieties (in particular those resistant to both viruses), or an unacceptable increase in costs (Legg et al. 2011; Patil et al. 2015; Rwegasira and Rey 2012). To understand which strategy is most likely to be successful, it is important to understand the decision-making process of a grower; what risks and costs are acceptable under what circumstances. Recent work has shown that this can have significant impact on the long-term success of disease control, and may represent the difference between success and failure (Carrasco et al. 2012; Legg et al. 2017; McQuaid et al. 2017a; Milne et al. 2015). At the same time, in order to attempt control on a regional scale, without which any local attempts at control will ultimately fail, it is important to understand how the viruses spread between fields and across distance. This is particularly relevant in the context of grower behaviour when considering the movement of planting material, which has been shown to be key in the spread of cassava viruses (Legg et al. 2014; Legg et al. 2011; McQuaid et al. 2017b; Patil et al. 2015).

Recently there have been a number of surveys assessing the impact and extent of CMD and CBSD in sub-Saharan Africa. Much of the work has concentrated on assessing the per-field disease incidence and severity on a regional level (Alicai et al. 2007; Chikoti et al. 2013; Gondwe et al. 2003; Hillocks et al. 1999, 2002; Mbewe et al. 2015; Mulenga et al. 2018; Rwegasira and Rey 2012). Conducted surveys were based on field observations, without assessing growers’ knowledge in terms of (i) capacity to identify the viral diseases of cassava, (ii) practices related to sourcing and exchange of planting material, and (iii) control strategies.

Although it has been shown that both CMD and CBSD pandemics depend strongly on the exchange of planting material (McQuaid et al. 2017b), and that growers often share their cuttings with friends and family (Houngue et al. 2018; Kombo et al. 2012; Ntawuruhunga et al. 2007; Teeken et al. 2018) there is a lack of studies that assess the distances over which this material is moved depending on sources or destinations. Thus, the primary objective of the current study was to obtain insight into the nature of the flow of cassava planting material into and out of the growers’ fields (specifically the volume moved over distance) depending on sources or destinations. The second objective was to ascertain grower knowledge of diseases, their symptoms and prevalence in the area, and the sources and preferences that growers had for obtaining this information. This information was gathered through a survey of growers across the country. The results of this work can be used to inform and improve disease control strategies, particularly those aimed at the recent outbreak of CBSD in Zambia. In particular, our investigation reveals the benefit and necessity of grower education programs, particularly through media and extension workers, to make growers active actors in the control of crop disease.

## Data and Methods

### Agro-ecological context of the study area

The study was conducted in five provinces of Zambia: Western, Luapula, Central, Northern and Eastern, which are among the major cassava growing areas and are known to have CMD. These provinces represent various environmental conditions (Figure 1). Northern and Luapula provinces are located in the Agro-Ecological Zone (AEZ) III, which comprises part of the Central African plateau and receives over 1000 mm of rainfall annually with a monomodal rainfall pattern (Saasa 2003; The World Bank 2006). The area has up to 190 days of growing season and is not prone to drought. The most widely practiced traditional farming systems by growers in this region are mainly based on “slash and burn” and shifting cultivation. The main crops grown include cassava, maize, sunflower, coffee, tea and many others (Ngoma 2008). Western, Central and Eastern provinces are located in a slightly drier AEZ II with a growing season of 120 to 150 days and receiving about 800 to 1000 mm of rainfall per annum (Jain 2007; The World Bank 2006). The farming system is mostly commercial and the major crops grown include maize, wheat, groundnuts and soy bean. In Luapula and the Northern province, the rainy season occurs between November and April, while in Eastern, Western and Central provinces the rainy season occurs between December and April. The rainy season is followed by a long dry spell lasting from May to October.

**Figure 1.**
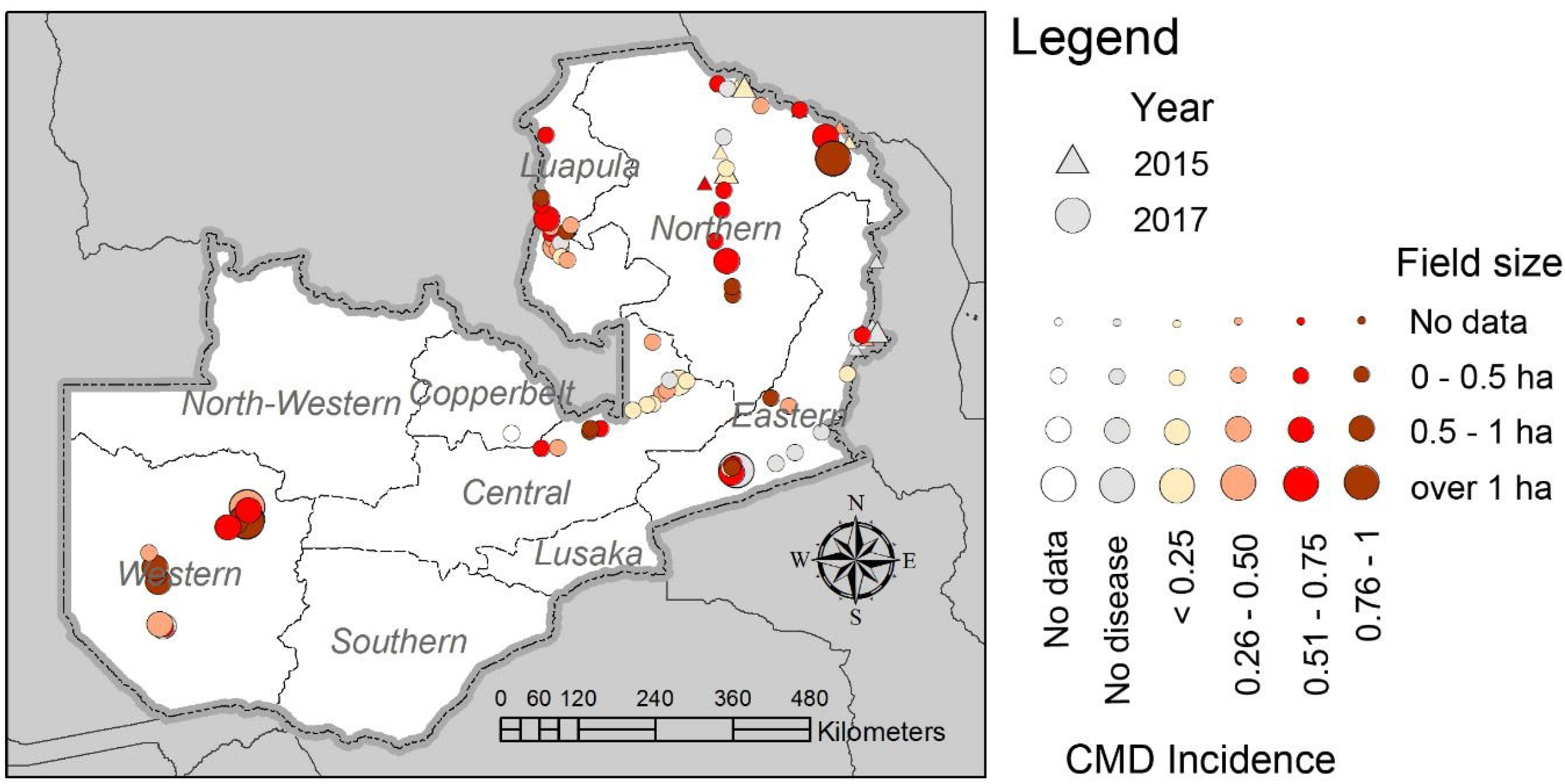
Locations of the interviewed growers in 2015 and 2017 in five provinces of Zambia, field sizes and CMD disease incidence (proportion of infected plants within the field).

### Sample selection

A total of 96 smallholder cassava growers were randomly selected along the major roads and were asked for permission before conducting the questionnaires and field samplings. 26 growers were interviewed in 2015 in the Eastern (10), Luapula (4) and Northern (12) provinces, and 72 growers were interviewed in 2017 in Central (15), Eastern (15), Luapula (15), Northern (14) and Western (13) provinces (Figure 1, Table 1). The research team comprised the principal investigator and two research assistants conversant with the local languages and with experience in cassava production for easy identification of the local varieties, CMD and CBSD symptoms. The study was conducted between January and May in both years. During this period, most plants were assumed to be between three to nine months old, as the rainy season generally starts in November in most parts of the country. Three to nine months after planting is generally regarded as being ideal for capturing foliar and root symptoms before the plants shed their leaves.

**Table 1.** Summary of the number and per-province distribution of the growers, average field size, variety number, planting frequencies and CMD incidence and severity scores. Incidence is calculated based on visual foliar symptoms. Mean severity scores are derived per field from symptomatic plants only. No visual CBSD symptoms were reported in the study.

| Province | Number of growers |  | Mean field size [ha] | Median number of varieties | Planting frequency |  |  | CMD |  |  |  |
| --- | --- | --- | --- | --- | --- | --- | --- | --- | --- | --- | --- |
|  |  |  |  |  |  |  |  | Incidence |  | Severity |  |
|  | 2015 | 2017 |  |  | Biennial | Yearly | Twice a year | Mean | SE | Mean | SE |
| **Central** | - | 15 | 0.26 | 2 | 0 | 2 | 2 | 36.7% | 7.6% | 2.93 | 0.189 |
| **Eastern** | 10 | 15 | 0.82 | 1 | 0 | 22 | 2 | 26.1% | 7.3% | 2.81 | 0.327 |
| **Luapula** | 4 | 15 | 0.29 | 3 | 1 | 18 | 0 | 47.9% | 6.0% | 2.74 | 0.156 |
| **Northern** | 12 | 14 | 0.45 | 2 | 0 | 23 | 0 | 43.8% | 6.5% | 2.82 | 0.142 |
| **Western** | - | 13 | 1.25 | 3 | 0 | 12 | 1 | 68.9% | 6.0% | 3.72 | 0.062 |

### Questionnaires

Structured interviews with a mix of closed- and open-ended questions were conducted with cassava growers who voluntarily agreed to participate (Szyniszewska et al. 2019). The questionnaire was pre-tested on a small group of growers before the survey and adjustments were made to ensure that the right information was obtained during the actual interviews. To encourage wider participation, the interviews and discussions were conducted in the local languages familiar to most growers in respective regions: Bemba for Northern, Luapula and Central provinces; Lozi for Western Province and Nyanja for Eastern Province. Some of the questions asked were repeated and rephrased to enable growers to understand and respond fully. The rephrasing was done without changing the original meaning of the questions.

In the first section of the survey general information on growers’ field location, altitude and field size was recorded. Growers were asked about varieties grown, planting and harvesting frequencies, and variety preferences and reasons. The second section of the survey comprised questions related to the trade of planting material: sourcing and exchange. Growers were asked how many bags went to or were obtained from their own fields, their stores, friends or family, markets, middlemen, NGOs or research stations, and how far away those sources or recipients were located. Growers were also asked about their favourite source of planting material and how frequently they use various sources. The third section of the surveys assessed growers’ knowledge of CMD and CBSD in terms of symptom recognition, presence of the diseases in their fields and surrounding areas, and the method of disease spread. The fourth and final section of the questionnaire was related to the sources and frequencies of obtaining information related to cassava diseases, certified clean seed systems (CSS) and the ranking of sources viewed as important to the grower. Growers were also asked about the factors that influence their decisions related to disease control, including disease pressure, their concern about the disease, and market prices that would encourage them to use CSSs.

### Disease incidence and severity

Plants at the fields visited were assessed for the presence and severity of disease foliar symptoms. In each field, a total of 30 plants were inspected, 15 plants on each diagonal line across the field (Sseruwagi et al., 20014). The plants were scored for the presence or absence of foliar symptoms of CMD and CBSD. Symptom severity for CMD was recorded on each plant using a five point rating scale (Hahn et al. 1980), where 1 = no disease symptoms; 2 = mild chlorotic pattern over entire leaflets or mild distortion at the base of leaflets only with the remainder of the leaflets appearing green and healthy; 3 = moderate mosaic pattern throughout the leaf, narrowing and distortion of the lower one-third of leaflets; 4 = severe mosaic, distortion of two thirds of the leaflets and general reduction of leaf size, and 5 = severe mosaic and/or distortion of the entire leaf and plant stunting. The presence or absence of CBSD symptoms on the leaves and stems was recorded for each plant using a scale of 1 to 5, where 1 = no apparent symptoms; 2 = slight leaf feathery chlorosis with no stem lesions; 3 = pronounced leaf feathery chlorosis, mild stem lesions and no dieback; 4 = severe leaf feathery chlorosis, severe stem lesions and no dieback, and 5 = defoliation, severe stem lesions and dieback (Gondwe et al. 2003).

### Data analysis

The grower’s responses together with disease incidence and symptom severity, were analysed using the R language for statistical computing (R Core Team 2016). Frequency distributions were plotted to illustrate and compare response rates for each category. Sets of descriptive statistics including means and standard errors and cross tabulations were calculated. Results were expressed as percentages of the frequency of responses obtained from growers, excluding records where data were not available (thus totals may differ in each question) and plotted with the *ggplot2* package (Wickham 2016). Logistic regression was used to relate grower’s disease knowledge with disease incidence using ‘glm’ function in the *lme4* package in R (Bates et al. 2015, p. 4) and χ^2^ contingency tests were performed using ‘chisq.test’ function.

## Results

### Field properties and varieties preferences

Most growers’ fields were small (mean 0.59 ha) and planted annually (92.9% of participants) (Table 1). Harvesting was based on need (40% of participants), from which we conclude that those surveyed were primarily small-scale subsistence growers. Incidence of CMD was generally high (range of 26.1 to 69.8%), while CBSD was not observed. Growers typically plant more than one variety of cassava in their fields (66.5% of visited locations). Good taste and associated sweetness (30 growers) and good yield and big tubers (21 growers) were the most commonly cited traits determining varietal choice (Supplementary Figure 1). The availability of planting material (14 growers) and early maturing (13 growers) were important to them. Among six preference criteria dictating choice of planting material (Figure 2) varietal preference was the highest ranked while availability related answers were ranked second and third.

**Figure 2.**
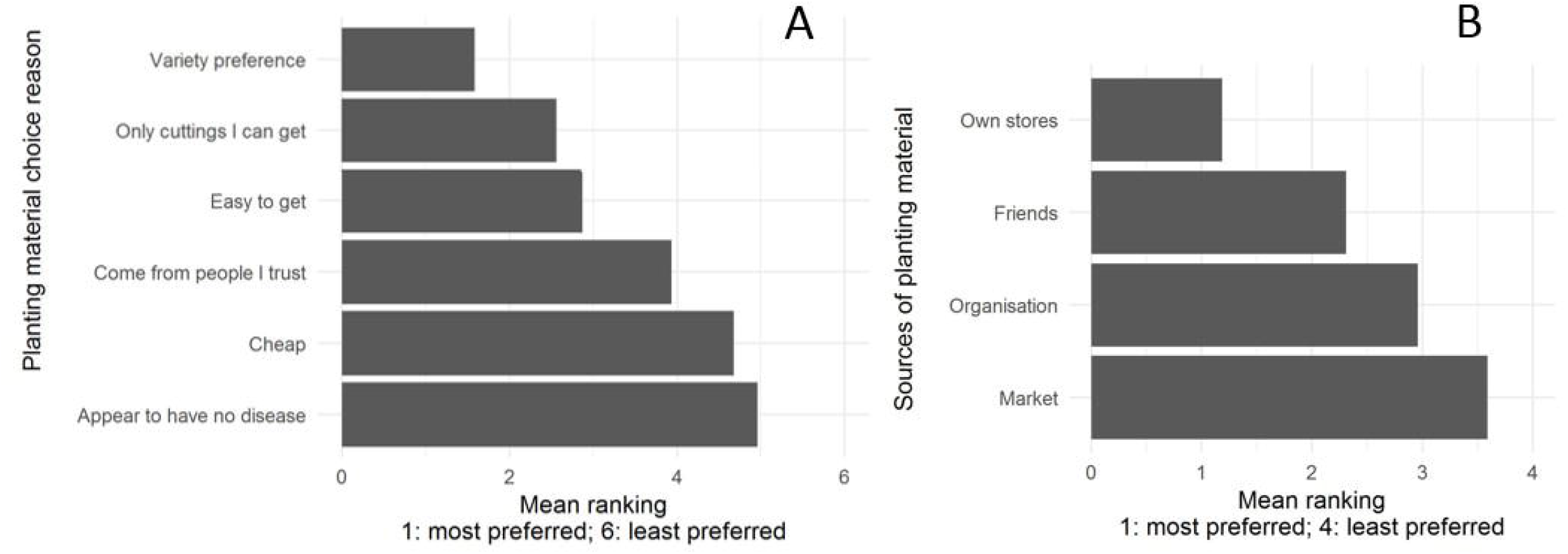
Planting material (A) choice reason and (B) preferred source. Ranking 1 represents most preferred.

### Planting material movement and trade

Most of the planting material was recycled from the previous crop (83 growers), stored (11 growers) or destroyed (52 growers) (Figure 3). While sharing did occur with family and friends (55 and 39 growers respectively) this was generally within the same or nearby villages with 94% of recipients located within a radius of 1-10 km, with a maximum of 100 km.

**Figure 3.**
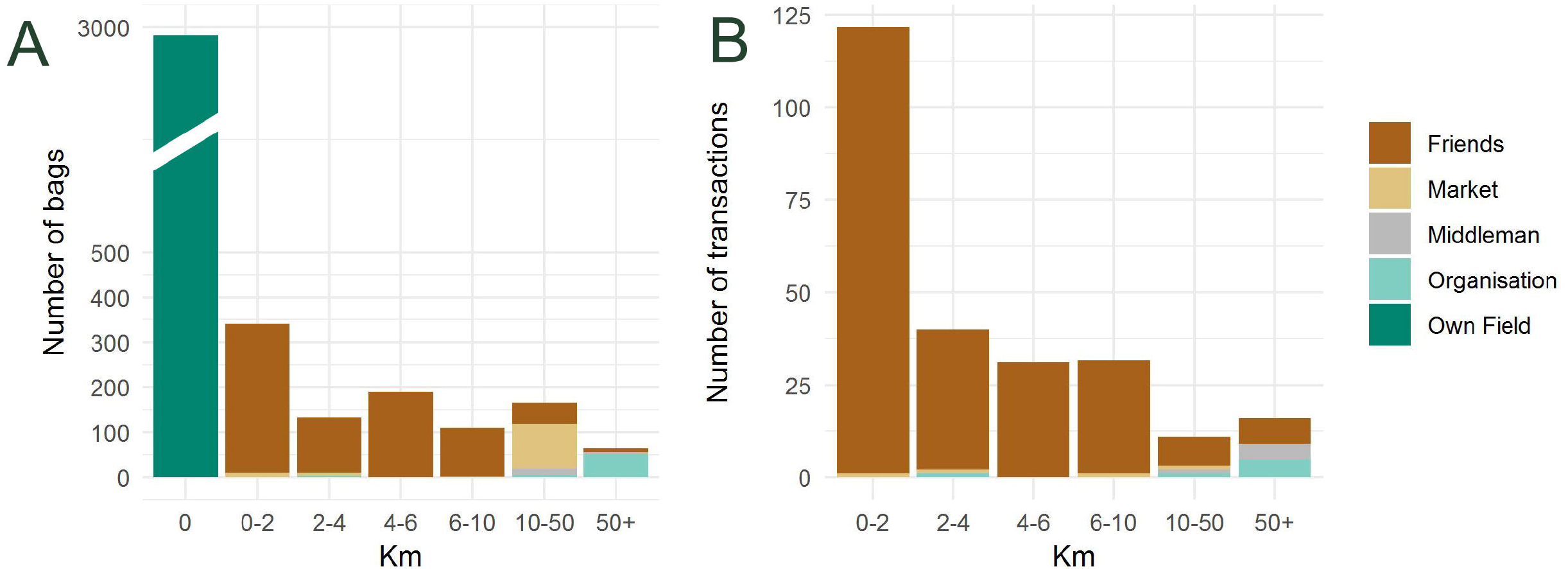
Total number of (A) bags of planting material moved and (B) individual transactions.

However, some movement did occur over a greater scale, including large transactions that moved planting material long distances to markets (100 bags over an average of 7.43 km), middlemen (9.5 bags on an average of 55 km), or NGOs (15 bags on an average of 28.5 km). Given the paucity of data on movement distances for cassava, we provide some additional detail on some individual transactions to illustrate the range of behaviours evident in a relatively small cohort. One transaction involved moving a large amount of planting material (100 bags) from a single grower, with a large field of 4 hectares and the distance to the market was 40 km. Three further transactions with the markets occurred. 10 bags sold at the market within a distance 0.05 km by a grower with a field size of 1.5 hectare. The remaining two transactions involved small purchases of planting material (7 and 1 bag respectively) by small-holder growers (field size up to 0.25 hectares) travelling 3 and 8 km to the market. Overall, the range of reported distances to the market was between 0–40 km. Growers, who obtained their planting material from middle-men, indicated transaction distances of 50 and 60 km. Six growers exchanged their planting material with an NGO or an organization with the distance range of 0–350 km.

### CMD and CBSD knowledge

Most of the growers surveyed (81%) had no knowledge of what CMD was. Having surmised it was a disease, most (60.5%) were unable to recognise it by its symptoms, or identify its means of dispersal (75.6%) or it’s likely effect on yield (39%). Higher CMD incidence in the field was a significant predictor of grower’s knowledge of disease in a logistic regression (p < 0.0001). Nearly half of the growers (44%) did not know whether the disease had an impact in their area, and another 44% observed disease impact on the crop. Of those that felt the impact of the disease, 25.9% identified lost yield. Disease incidence did not prove to be a significant predictor of the answer whether or not the disease had an impact in the area.

Overall, when asked how concerned they were about CMD on a scale from 1 (least worried) to 10 (very worried), 53% of growers responded they had very low levels of worry (1-3), 17% of growers were moderately worried (4-6) and 28% were extremely worried (7-10). When we grouped them by how informed they were, growers with no information were less concerned compared with those that were informed (Figure 4).

**Figure 4.**
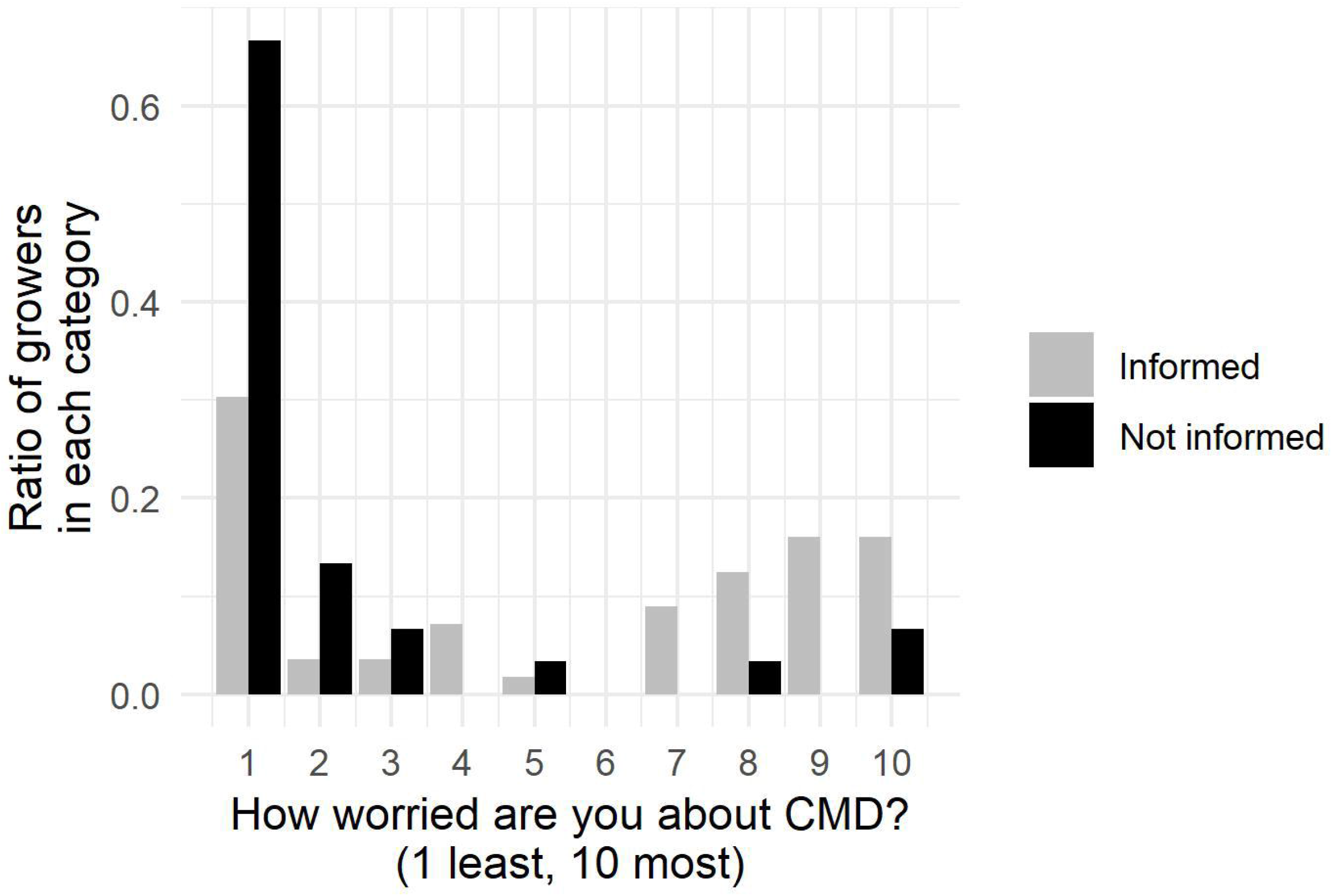
Growers response to the question: How worried are you about cassava mosaic disease, on a scale of 1 to 10 where 1 is the least worried and 10 is the most worried. Growers are categorised based on whether they had access to information about CMD in the past at least on one occasion (informed) vs those who did not have access to information about CMD (not informed).

None of the growers had knowledge about CBSD and no disease symptoms were detected in the field surveys.

### Disease control and management

Disease management for CMD is rare among the growers. Two thirds of the growers (74.7%) declared that they do not institute any control measures. In contrast, of the few growers that applied control measures, we found that five used clean planting material; two growers who were seeking help from agricultural officers, rouged the diseased plants, and sprayed for insects. The majority of the growers who used control measures were located in the Eastern province (8 out of 11). Most growers who implemented disease management cited their own experience as a source of planting knowledge (7), two cited agricultural extension officers and one grower cited parents and one a cooperative group.

### Certified clean seed (CCS) sourcing and knowledge

Nearly half of the growers interviewed (46%) were aware of CCS, and nearly half of them would seek it from agricultural extension workers. At the same time, of those who were unaware of CCS, the majority (28%) would be happy to use them if available, and no growers indicated that they would not be happy to use CSS if it were provided or available.

### Information sources

Among the surveyed growers, 21.4% identified agricultural extension workers as a source of information, while 30% relied on information on cassava planting practices passed on from their parents and grandparents and 27.4% relied on their own experience in farming as their source of knowledge. Information on cassava diseases and CCS reached half of the growers on at least one occasion (50.6% and 51.8% respectively), although no single source of information reached the majority of individuals. The most frequent sources of information included nearby friends, family and neighbours, and the radio.

In terms of growers’ preferences for information, extension workers, radio and people within the village were clearly favoured (Figure 5), while the village leader or distant friends or relatives were least preferred. Nearly 90% of growers who were aware of CMD had access to frequent information, whilst the majority of growers who were unaware of the disease had no access to information (Figure 6). Most informed growers were located within the Northern and Eastern provinces, where over half of growers had often heard about CMD from various information sources. The least informed growers were located in Luapula and Western provinces, where over two thirds of the growers reported never receiving information about CMD.

**Figure 5.**
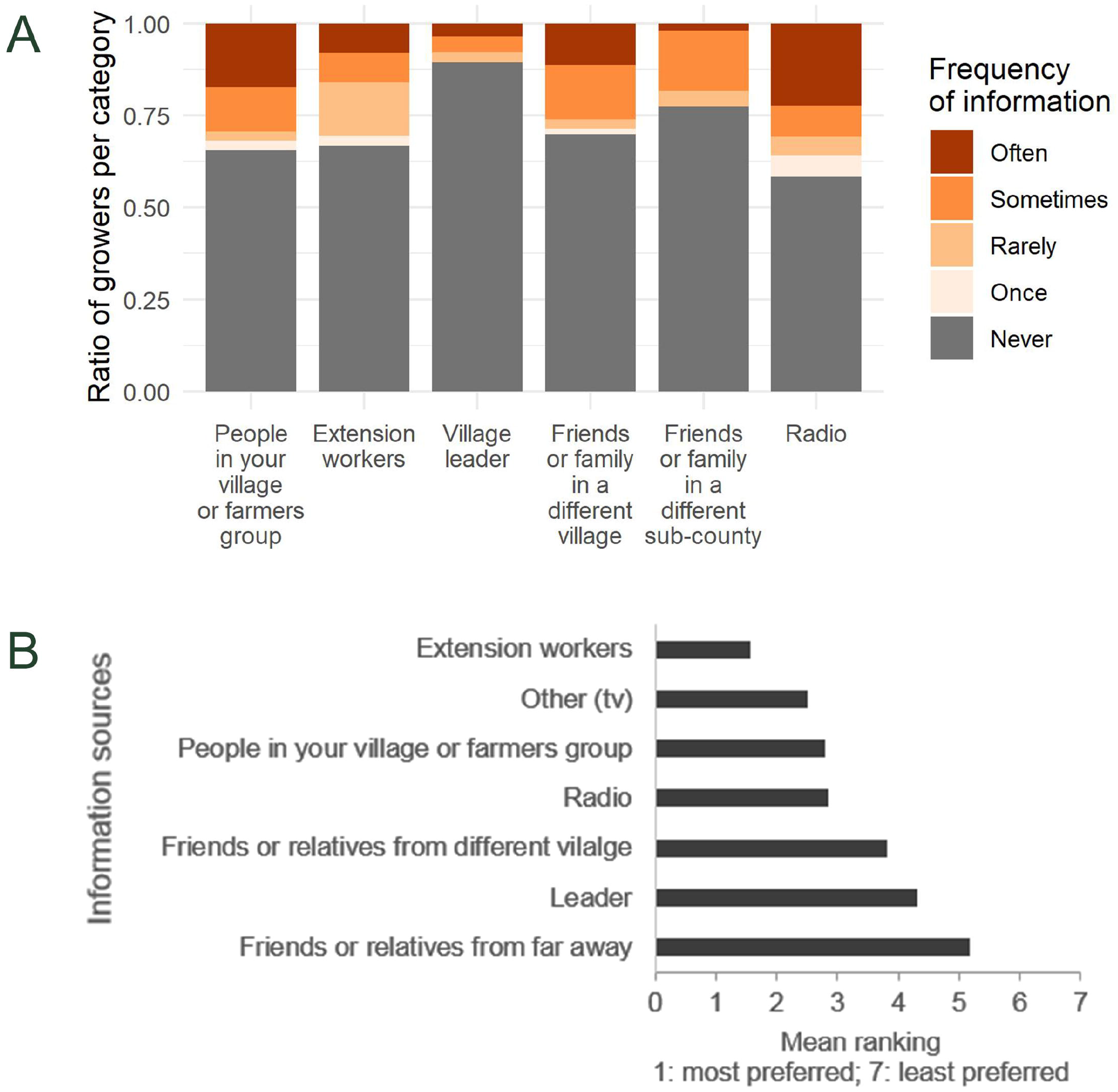
(A) Frequency of hearing information about cassava diseases from various sources and (B) ranking of information sources.

**Figure 6.**
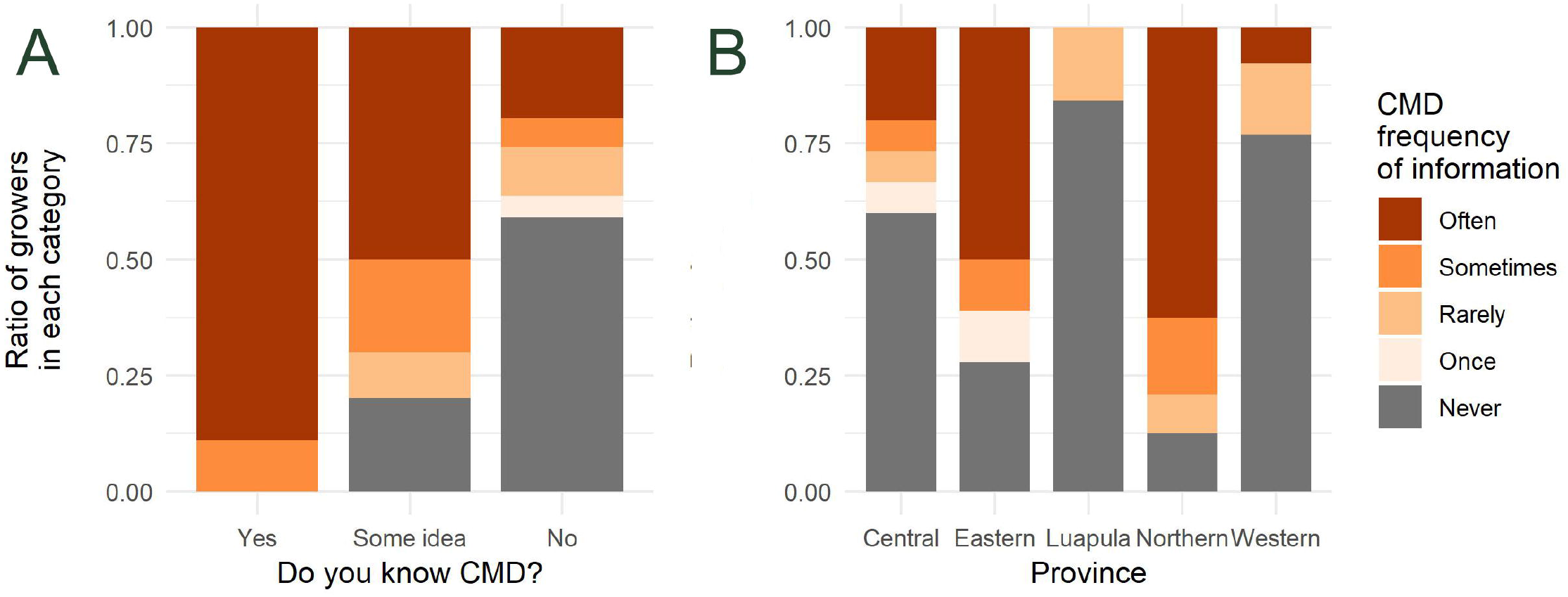
Disease knowledge vs frequency of obtained information (A) and by province (B).

### Making decisions

High yield, cost and lack of disease were the most frequently reported factors (27.4%, 25% and 22.6%, respectively) that would influence growers’ decision on whether or not to use certified clean planting material. Surprisingly, few growers (3.6%) would consider adoption of CCS if it were free. Majority of the interviewed growers indicated they would consider adoption of the CCS or would control for CMD if two to four neighbours would be affected or use it (Supplementary Figure 2).

Growers were classified as having knowledge, some knowledge or no knowledge. In those three categories 40%, 18% and 8% of growers respectively controlled for the disease. However, differences between these groups were not statistically significant (χ^2^ test *p* = 0.19). The price of clean seed did play a role in decision-making, with the intention to buy clean seed linearly decreasing with increasing price.

## Discussion

Cassava virus diseases constitute a major constraint to the production of cassava in sub-Saharan Africa, yet there have been few studies looking into one of the key aspects of disease spread or control; the knowledge and decision-making of the cassava growers themselves. It is widely acknowledged that the burden of these diseases can be amplified within an individual field by replanting infected material (Samura et al. 2017), and on a larger scale by sharing planting material between fields (McQuaid et al. 2017a; McQuaid et al. 2017b; Patil et al. 2015), yet there are no studies of which the authors are aware that investigate the physical properties of human mediated transmission. This is critical from the point of view of disease management and control, in particular for a complete understanding of disease spread that underpins effective disease management.

According to our survey, cassava seed trade is largely informal in Zambia, except for a limited number of commercial growers involved in the production and sale of planting materials. Growers mostly recycle materials from their own fields, attributing this to variety preference as well as the fact that the material is readily available. The preference for recycling is supported by previous studies, which have shown that a majority of planting material is recycled within the same field, while a considerable portion is also exchanged with close friends or family (Chikoti et al. 2016; Gnonlonfin et al. 2011; Houngue et al. 2018; Ntawuruhunga et al. 2007; Teeken et al. 2018). Although markets, organisations and middle-men are rarely involved in the movement of planting material, the large scale of the distances and quantities of material moved in each of these transactions highlighted by our study does indicate that they could be responsible for the movement of disease across large distances, which previous work has demonstrated could be severely detrimental to disease control (Legg et al. 2014; McQuaid et al. 2017a; McQuaid et al. 2017b). Increased trade movement of infected planting material could increase its importance in dispersal of CMD and CBSD still further (McQuaid et al. 2017b).

In general, most growers indicated that markets were more than 7 km from their homesteads. As presented in the study of Salasya et al. (2007), the closer a household is to the market, the higher the probability of adoption of improved varieties by that household due to greater market accessibility. Growers further away from markets are at a disadvantage, as they may lack market information and thus be more inclined to subsistence production. As a result, they may be less interested in the use of improved varieties as long as traditional varieties provide subsistence for the family. Growers are also, of course, sensitive to the price of planting material, and an increase in the price of improved seed relative to the local variety will reduce the adoption rate (Langyintuo and Mekuria 2008). From our study, however, it appears more likely that a lack of knowledge and access is a more significant hindrance, which must be considered when implementing clean seed systems.

Our work supports numerous previous studies that have shown that culinary properties and taste of planting material is as important, if not more important, in planting material selection than properties of more economic traits such as yield, while the appearance of disease makes little to no difference on choice (Houngue et al. 2018; Kombo et al. 2012; Njukwe et al. 2013; Ntawuruhunga et al. 2007). With this in mind, efforts to use clean (and possibly also disease-resistant or tolerant) planting material to control disease epidemics need to address growers’ varietal preferences and needs (Evenson and Gollin 2003; Kiros-Meles and Abang 2008). If new varieties are not suited to local tastes the level of adoption is likely to be low, a factor to be considered by both cassava breeders and clean planting material producers alike. At the same time, the importance of yield in varietal choice presents an opportunity to educate and reassure growers about the economic advantages of clean seed systems and the adoption of improved varieties.

Indeed, the lack of attention given by growers to the appearance of disease on a plant, or the decision to try and control for the disease, appears to be primarily due to a striking lack of knowledge about disease despite its widespread prevalence in growers’ fields. While this is unsurprising for CBSD, CMD has been present across the country for more than two decades, incurring estimated yield losses of between 50 – 70% (Muimba-Kankolongo et al. 1997). This is a reflection of the scarcity of information available to growers; only half of growers receive any information on disease or its control at some point, and few receive information frequently or on a regular basis. Access to information is critical towards decision making, and increases concern about disease impact at the very least, as our results show.

Lack of awareness about the risk and impact of disease on yield in turn could lead to the failure of disease control measures implemented at a wider level, where it is necessary for a large proportion of growers to engage in disease management in order for effective, sustainable control to work (McQuaid et al. 2017a). It is certainly highly likely that the lack of awareness, combined with high incidence, likely contributes significantly to the spread of the disease. At the same time, the high use of growers’ own planting materials, due to a lack of alternative sources, likely results in material susceptible to pests and diseases with a low genetic potential - similar observations have been made in Malawi (Chipeta et al. 2016).

The results underscore the important of role of extension workers in providing information to growers. Regular visits of trusted extension workers are required to provide growers with information on innovation, general crop production, marketing and disease control strategies. Although in our study extension workers were the most trusted source of information, only a small proportion of growers were reached by this means. Our results demonstrate the need for grower education to improve knowledge and create awareness that is vital in controlling disease. Although other sources of information, such as radio, TV, or mobile phone apps can certainly be helpful in reaching growers and should not be ignored, to bridge the gap between scientific and indigenous knowledge, substantial effort should be invested in extension workers to train growers in disease recognition, the impact of the disease, and the means of spread and, most importantly, control. Reducing the presence of cassava virus diseases, and increasing the yields of small-holder growers across Zambia and East Africa, will not happen without well-informed growers acting at the individual level.

## Conclusion

We have shown for the first time how far and how much cassava planting material moves due to trade. It appears that trade is likely responsible for much of the spread of viral diseases, where growers are unaware of this effect, as well as the disease itself, and consequently do little to prevent it. Elsewhere we see that grower awareness and education can be vital to engagement with disease control measures, so this lack of awareness highlights the need for grower education. The optimal manner in which to achieve this is through widely-trusted extension workers, although a number of other avenues such as farmers’ groups and radio are also important.

In conclusion, in order to control cassava virus diseases, we need clean seed systems and improved (resistant or tolerant) varieties. For these to be effective, growers need to be educated in the diseases, and to achieve this we need to utilise and strengthen the existing extension worker network as well as make use of farmer’s groups and the media.

## Acknowledgments

A.M.S., C.F.M., C.A.G. and F.v.d.B. gratefully acknowledge funding from the Bill and Melinda Gates Foundation grant to the University of Cambridge. Rothamsted Research receives support from the Biotechnology and Biological Sciences Research Council, United Kingdom.

P.C.C., M.T and R.M. are grateful to Mikocheni Agricultural Research Institute for the financial assistance through grant number OPP1052391 with funds provided by the Bill and Melinda Gates Foundation (BMGF). All authors are indebted to all the participating farmers for providing responses and allowing surveyors to access their cassava fields.

## Authors contributions

C.F.M. and F.v.d.B conceived the study, designed the questionnaire and drafted the manuscript. A.M.S. analysed the data and drafted the manuscript. P.C.C. led the fieldwork and drafted the manuscript. M.T and R.M. carried out the fieldwork and drafted the manuscript. C.A.G. drafted the manuscript. All authors gave final approval for publication.

## Conflict of interest

The authors declared that they have no conflict of interest

---

**Supplementary Figure 1.**
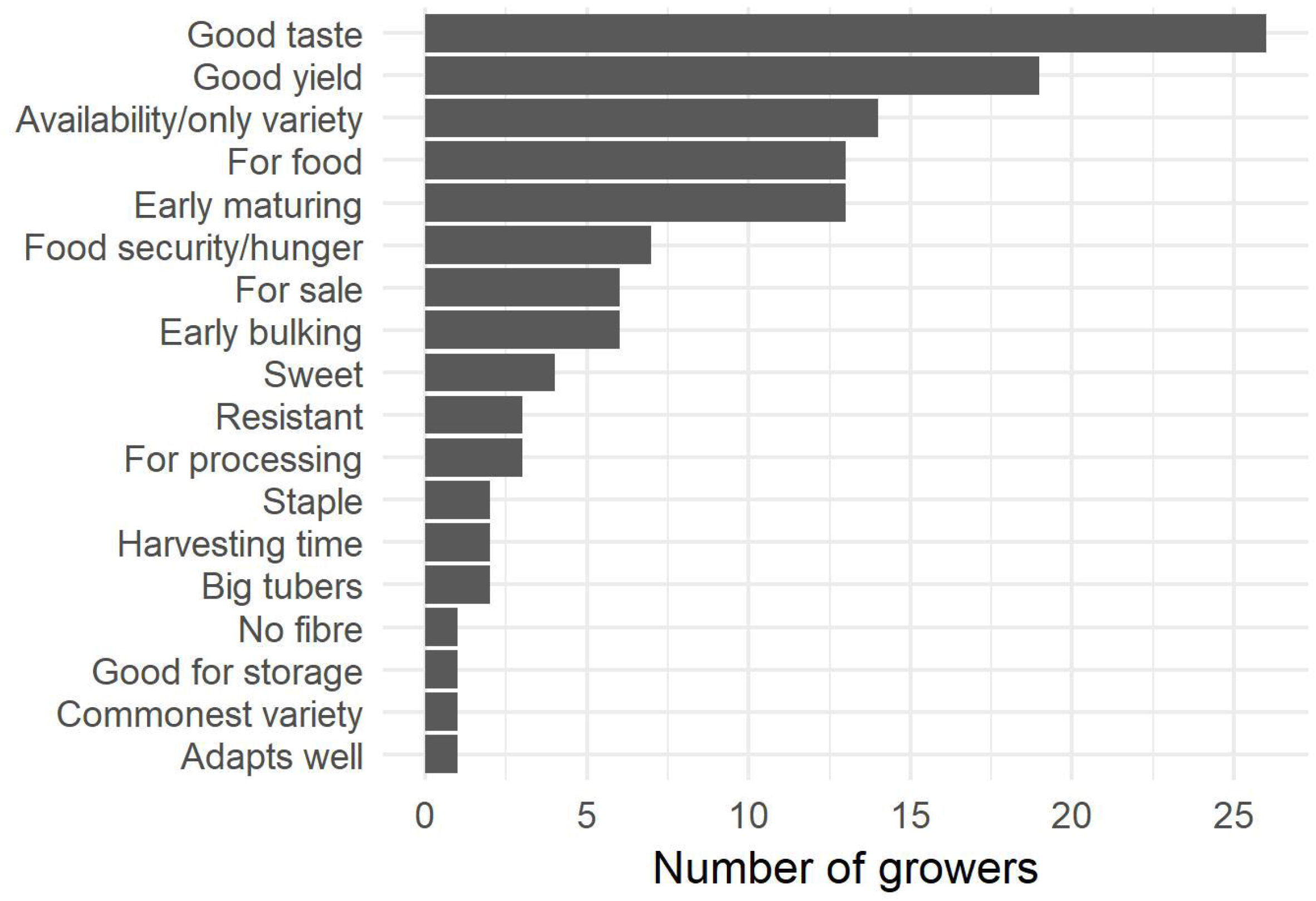
Different cassava traits dictating varietal choice cited by growers.

**Supplementary Figure 2.**
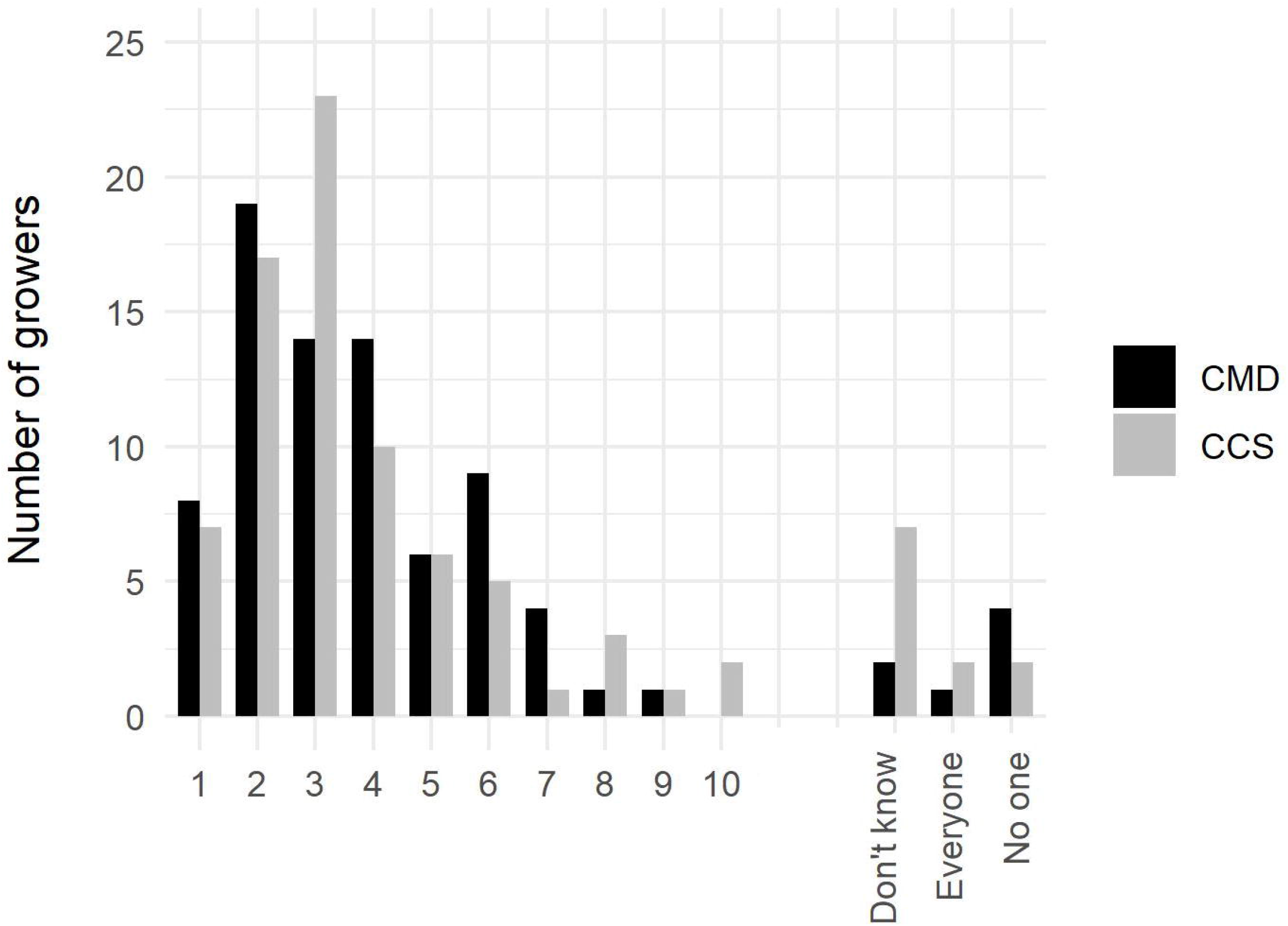
Response to the question: After how many neighbours have the disease (CMD) or use certified clean seed (CCS) would you think about control?

